# Metabolic potential of uncultured Antarctic soil bacteria revealed through long-read metagenomic sequencing

**DOI:** 10.1101/2020.12.09.416412

**Authors:** Valentin Waschulin, Chiara Borsetto, Robert James, Kevin K. Newsham, Stefano Donadio, Christophe Corre, Elizabeth Wellington

## Abstract

The growing problem of antibiotic resistance has led to the exploration of uncultured bacteria as potential sources of new antimicrobials. PCR amplicon analyses and short-read sequencing studies of samples from different environments have reported evidence of high biosynthetic gene cluster (BGC) diversity in metagenomes. However, few complete BGCs from uncultivated bacteria have been recovered, making assessment of BGC diversity difficult. Here, long-read sequencing and genome mining were used to recover >1400 mostly complete BGCs that demonstrate the rich diversity of BGCs from uncultivated lineages present in soil from Mars Oasis, Antarctica. The phyla Acidobacteriota, Verrucomicrobiota and Gemmatimonadota, but also the actinobacterial classes Acidimicrobiia, Thermoleophilia, and the gammaproteobacterial order UBA7966, were found to encode a large number of highly divergent BGCs. Our findings underline the biosynthetic potential of underexplored phyla as well as unexplored lineages within seemingly well-studied producer phyla. They also showcase long-read metagenomic sequencing as a promising way to access the untapped reservoir of specialised metabolites of the uncultured majority of microbes.

## Introduction

Throughout the last century, bacterial natural products have proven invaluable for humankind. Their diversity has been harnessed to treat different ailments, and above all, to fight infectious disease. However, their biological roles and even the extent of their diversity are not well understood. Over the last decade, metagenomics has shown that a vast amount of the bacterial diversity on Earth is comprised of uncultured bacterial taxa, with 97.9% of bacterial operational taxonomic units (OTUs) estimated as unsequenced^1^. First efforts to characterise and harness the specialised metabolite diversity encoded in metagenomes have shown promising results^2–4^. Metagenomic library screenings have yielded novel compounds, among them antibiotics^3,5,6^, while sequence-based studies have documented their diversity. In a study of grasslands with 1.3 Tb of short-read sequence data, Crits-Cristof et al. recovered hundreds of metagenome-assembled genomes (MAGs) obtained through a combination of binning approaches^7^. Analysis of the MAGs revealed a large number of BGCs in Acidobacteria and Verrucomicrobia, widespread but underexplored phyla of soil bacteria. Analysis of nonribosomal peptide synthetase (NRPS) and polyketide synthase (PKS) domains indicated that NRPS and PKS from these groups were highly divergent from known BGCs of these classes. Borsetto et al. also reported a high degree of diversity of NRPS and PKS domains in Verrucomicrobia and other difficult-to-culture phyla^8^. Finding efficient ways to access this treasure trove of diverse and unexplored specialised metabolites will expand our understanding of microbial natural products, yield novel and useful compounds, and be an important step towards the development of much-needed antimicrobials.

Recent advances in long-read sequencing technology have made it possible to recover largely complete genomes metagenomic sequencing projects. A sequencing effort of 26 Gb returned 20 circular genomes from human stool samples^9^, while a recent study using 1 Tb of long-read data from wastewater treatment plants recovered thousands of high-quality MAGs, 50 of which were circular^10^. Using mock community data, Pérez et al. demonstrated that full-sized BGCs could be successfully recovered from long-read metagenomic sequencing^11^.

In recent years, a number of tools to explore and understand BGC diversity have been developed. Genomes can be mined for known classes of BGCs using tools such as antiSMASH^12^, while the MiBiG database^13^ links BGCs to known compounds. BGCs can then be compared in networking-based tools such as BiG-SCAPE^14^ and BiG-SLiCE^15^ to assess relations of BGCs and estimate their novelty relative to extant BGC databases.

The isolated, harsh and unique environments of Antarctica show high degrees of endemism in their bacterial life, but their diversity remains underexplored^16^. Little is known about the specialised metabolites of Antarctic microorganisms. Few studies have explored polar, and specifically Antarctic, natural products using functional screening of isolates and metabolomics^17–21^. A high number pigmented bacterial isolates indicates that carotenoids and PKS, among other pigments, could be abundant BGC classes^22^. One culturing study suggested that Antarctic isolates show a below average potential for antimicrobials^17^. On the other hand, a primer-based study showed a promising diversity of NRPS and PKS diversity in soil from Mars Oasis in the southern maritime Antarctic^8^, a site with exceptionally high diversity of micro- and macroorganismal life for its latitude^23,24^. Low-temperature, aerated Antarctic soils have previously also been linked to methanotrophy^25,26^, and these soils could therefore harbour methanobactins, small ribosomally synthetised peptides that scavenge copper needed for methane monooxygenases.

In the present study, we used long-read shotgun metagenomic sequencing coupled with genome mining and bin- and contig-based taxonomic classification to analyse the biosynthetic potential of soil from Mars Oasis. We recovered >1,400 highly diverse and mostly complete BGCs from largely uncultured and underexplored bacterial phyla such as Acidobacteriota, Verrucomicrobiota and Gemmatimonadota as well as hitherto uncultured members of Proteobacteria and Actinobacteriota. This helps elucidate the biosynthetic diversity and highlights potential applications of the underexplored Antarctic soil microbiome. The present study further demonstrates how long reads make BGC recovery, analysis and taxonomic classification from highly complex metagenomes feasible even at low sequencing efforts (<100 Gb).

## Materials and Methods

### Site description

Mars Oasis is situated on the south-eastern coast of Alexander Island in the southern maritime Antarctic at 71° 52’ 42” S, 68° 15’ 00” W (Figure 1A). Mean soil pH is 7.9, with NO^3-^-N and NH^4+^-N concentrations of 0.007 mg kg^-1^ and 0.095 mg kg^-1^, and total organic C, N, phosphorus and potassium concentrations of 0.26%, 0.02%, 8.01% and 0.22%, respectively. Soil moisture concentrations range between 2% and 6% in December–February, when snow or rainfall events are very rare, with the majority of precipitation falling as snow between March and November. Mars Oasis has a continental Antarctic climate, with frequent periods of cloudless skies during summer, when temperatures at soil surfaces reach 19 °C. During midwinter, the temperatures of surface soils decline to -32 °C. Mean annual air temperature is *c*. -10 °C^27^.

**Figure 1:**
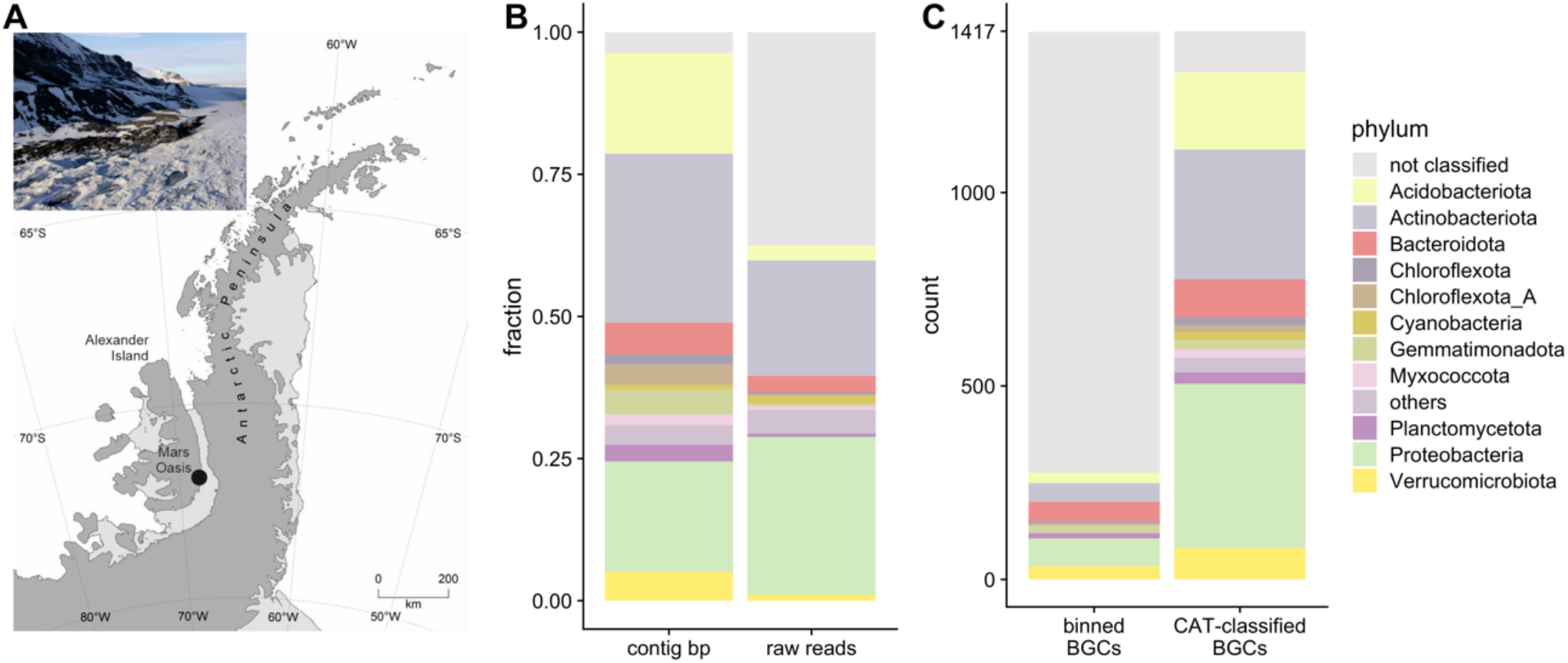
(A): Map of the Antarctic Peninsula with Mars Oasis indicated. Inset: Aerial photo of the site taken in austral summer; (B): Phylogenetic classification of contigs (by CAT) and reads (by kraken2); (C) phylogenetic classification success of BGCs from binned contigs and CAT-classified contigs.

### Soil sample, extraction and sequencing

Four samples of surface soil (each *c*. 2.5 kg) were collected from the lower terrace at Mars Oasis by British Antarctic Survey staff in 2018 and were kept cool for several hours before being stored at -20 °C. Soils were kept at this temperature until DNA extraction. A gentle chemical lysis and DNA extraction were performed and the DNA was subjected to size selection to approximately 20 Kb and larger by agarose gel electrophoresis using a protocol previously used for metagenomic library construction^28^. DNA was sequenced using Oxford Nanopore Technologies (ONT) MinION and Illumina HiSeq 150 bp paired-end reads. For long reads, the DNA was sequenced using three R9.4.1 flow cells and the SQK-LSK109 kit. The nuclease flush protocol was used between each independent library run on a flow cell. Short read DNA library preparation and Illumina sequencing were performed by Novogene according to their in-house pipeline. In short, one µg of DNA was sheared to 350 bp, then prepared for sequencing using NEBNext® DNA Library Prep Kit. The library was enriched by PCR and underwent SPRI-bead purification prior to sequencing on a HiSeq sequencing platform.

### Assembly, polishing and quality control

The long reads raw data were basecalled with Guppy v.3.03 (HAC model) and assembled using Flye^29^ v2.5 using the --meta flag. The resulting assembly was polished with 4 iterations of Racon^30^ v1.4.7 followed by one run of Medaka^31^ v0.7.1. Then, the short reads were used for six rounds of polishing with pilon^32^ v1.23. The approximate assembly quality was checked at every step using ideel^33^. Read and assembly statistics can be found in Results Table 1. Initial assessment of potential indels showed that 82% of all proteins were shorter than 0.9 times the length of the closest reference protein in the UniProt database and 7.2% were longer than 1.1 times the length of the closest reference protein. After polishing using Racon, Medaka and pilon, the proportion of potentially truncated proteins was reduced to 70%, while that of proteins that were potentially too long slightly increased to 7.6%.

**Table 1:**
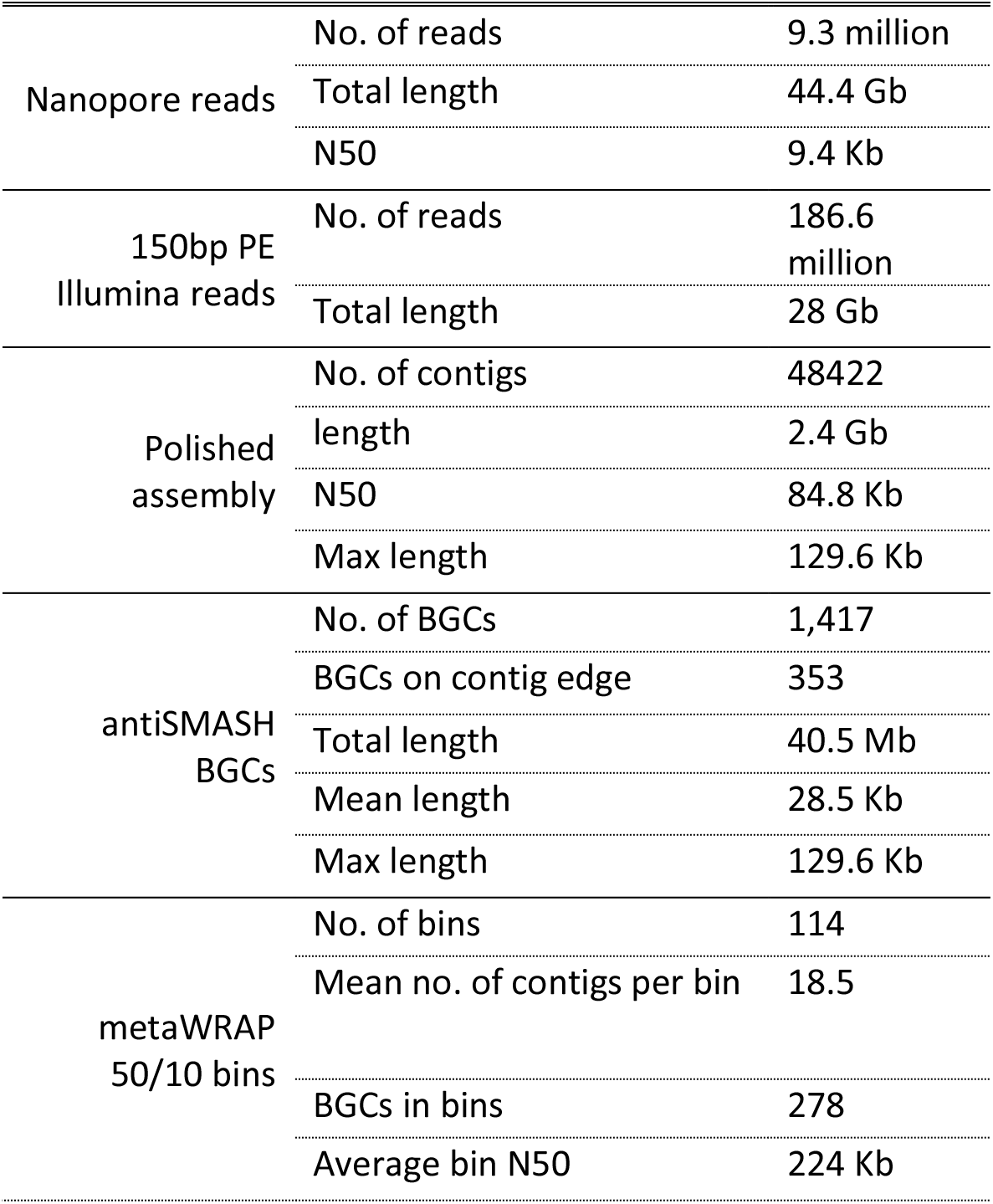
Raw sequence, polished assembly, BGC mining and binning statistics

### Genome mining, binning, taxonomic assignment and quality control

For detecting biosynthetic gene clusters, the polished assembly was analysed by antiSMASH^12^v5.1. For taxonomic assignment of contigs, proteins were predicted using Prodigal^34^, and CAT^35^ (settings --sensitive -r 0.5 and -f 0.3) was used with a DIAMOND^36^ database built from proteins in the GTDB_r89_54k database^37^ as well as the NCBI non-redundant protein database. The contigs were also binned with MetaBAT2^38^, CONCOCT^39^ and MaxBin2^40^, using long- and short-read abundance profiles for differential coverage. The resulting bins were subjected to metawrap-refine^41^ to produce the final bins. BiG-SCAPE^14^ 1.0.1 was run in --auto mode with --mibig enabled to identify BGCs families. Networks using similarity thresholds of 0.1, 0.3, 0.5 and 0.7 were examined, since higher thresholds led to extensively large proposed BGC families. In order to calculate BGC novelty, BiG-SLiCE 1.1.0^15^ was run in --query mode with a previously prepared dataset which had been computed from 1.2 million BGCs using --complete_only and t = 900 as threshold^42^. The resulting distance *d* indicates how closely a given BGC is related to previously computed gene cluster families (GCFs), with a higher *d* indicating higher novelty. For this analysis, we highlighted values of *d* > t and *d* > 2t (i.e. *d* > 900 and *d* > 1800, respectively), as they were previously suggested as arbitrary cutoffs for “core”, “putative” and “orphan” BGCs^42^.

### Precursor peptide homology searches and sequence logo construction

ORFs were aligned using Clustal Omega^43^ and a HMMER^44^ search was performed in the EBI reference proteome database with a cut-off E-value of 10E-10. The resulting protein sequences were aligned using Clustal Omega and a HMM was generated and visualised using skylign.org^45^.

## Results

### Taxonomic classification and binning of BGCs

Contigs were binned using CONCOCT, MaxBin2 and MetaBAT2, and consensus bins were generated using metaWRAP refine. This yielded 114 bacterial bins with CheckM completeness > 50% and contamination < 10% containing 278 BGCs (see Table 1.) Since only 278 BGCs had been binned, a contig-based classification approach was adopted. All contigs were classified using CAT with a database based on Genome Taxonomy Database (GTDB) r89 proteins, leading to a classification of 93% of BGC-containing contigs at a phylum level (Figure 1B-C). A cross-check of bin-level classification and contig-level classification of BGC-containing contigs showed no conflicting assignments. Of the 2,980 total binned contigs, 71 (2.4%) were classified differently at order level using CAT. Bin-level classification was preferred where available.

### Recovery of diverse and complete BGCs

The polished assembly was analysed using antiSMASH v5.1. A total of 1,417 BGCs were identified on 1,350 contigs (Table 1). A total of 353 BGCs (24.9%) were identified as being on a contig edge and were therefore categorised potentially incomplete. The most abundant classes of BGCs were terpenes (27.2%), followed by NRPS (15.7%) and bacteriocins (10.1%). In particular, terpenes were dominated by few subclasses. Out of 401 observed terpene BGCs, 321 contained a squalene/phytoene synthase Pfam domain (PF00494). This indicates that the product of these BGCs is a tri- or tetraterpene. Forty-four BGCs also contained a squalene/hopene cyclase (N terminal; PF13249), 39 BGCs contained a carotenoid synthase (PF04240), while 47 contained a lycopene cyclase domain (PF05834).

Approximately half of the ribosomally synthesized and post-translationally modified peptides (RiPPs) identified in the sample contained methanobactin-like DUF692 domains (PF05114). However, no BGCs resembling known methanobactin BGCs were found.

The proportion of proteins identified as too short on BGC-containing contigs was estimated at 63%. It is possible that this measure was influenced by the UniProt reference database not containing representative proteins for the mostly uncultivated strains recovered in this study. However, fragmentation of ORFs through indels was clearly visible, especially in NRPS and PKS BGCs in which whole megasynthase genes were broken up into several fragments.

### Long reads and GTDB improve phylogenetic classification of environmental BGCs

The use of GTDB proteins instead of the NCBI non-redundant protein database increased the classification success of BGC-containing contigs from 36.8% classified at order level with the NCBI database to 71.8% with GTDB. The difference was mainly due to BGCs from MAG-derived orders which were not present in the NCBI database, such as UBA7966. However, the GTDB database is also much smaller than the NCBI nr database, and many MAG-derived clades especially at lower taxonomic ranks do not have many representatives in the GTDB database. To avoid misclassifications, we therefore decided to conduct analysis at class and order level, even if contigs were classified at lower taxonomic ranks.

To assess the advantages of long-read sequencing for BGCs detection and classification, the output was compared with Biosyntheticspades, which allows the assembly of NRPS and PKS from short-read sequences by following an ambiguous assembly graph using *a priori* information about their modularity. Using Biosyntheticspades with the 28 Gb of short reads, 228 unambiguous NRPS/PKS BGCs were predicted. Sixty-one of these were above 5 Kb long and five NRPS were larger than 30 Kb. Furthermore, 202 other BGCs were predicted from other contigs. Classification success with CAT using GTDB was comparatively lower, with only 70% classified at phylum level, and 54% classified at order level. This could be attributed to the fact that Biosyntheticspades does not assemble the genomic context around the BGCs. The phylogenetic classification of BGCs reflected the composition found using the nanopore assembly. While Biosyntheticspades predicted a large number of BGCs in total, the practical usability and interpretability of the output remained low, since completeness, cluster borders and potential modification genes could not be assessed and phylogenetic classification success was reduced.

### Highly divergent BGCs found in unusual specialised metabolite producer phyla

Examination of the BGC counts by BGC type and phylum showed that the three well-known producer phyla Actinobacteriota, Proteobacteria and Bacteroidota together contributed over 60% of BGCs (Figure 2A). BGCs attributed to Acidobacteriota and Verrucomicrobiota represented up to 20% of the total BGCs, while other phyla constituted the remaining 12%, and 7% remained unclassified at phylum level. In particular, 20% of NRPS remained unclassified at phylum level. No archaeal BGCs were found.

**Figure 2:**
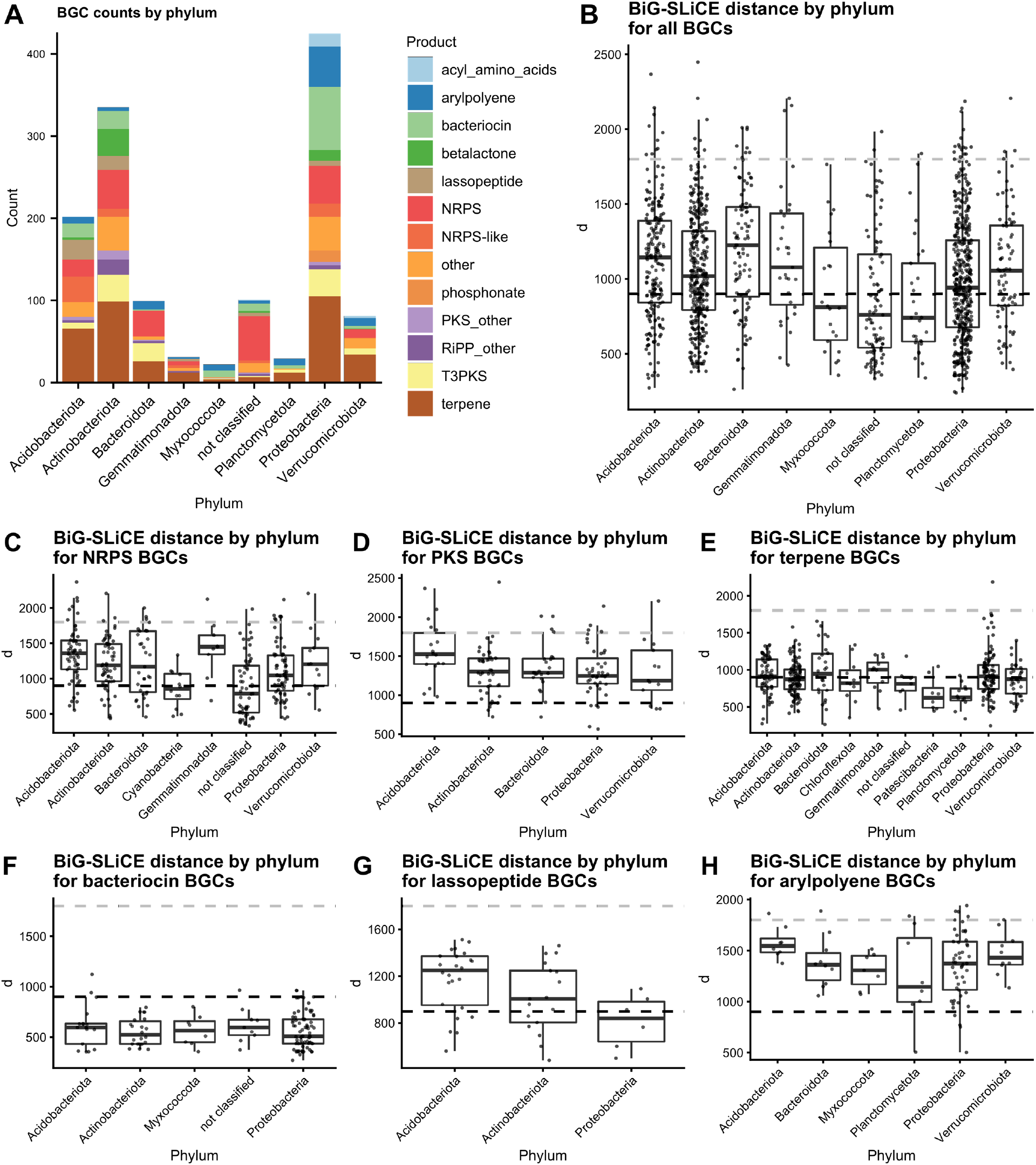
(A) BGCs by phylum and BGC type (phyla with a count <20 removed; products with count <10 under “others”, (B) BiG-SLiCE distances of BGCs by phylum, with the black dotted line indicating d = 900 and the grey dotted line d = 1800 (phyla with a count <20 removed); (C-H) BiG-SLiCE distances for different BGC types plotted by phylum (phyla with < 5 BGCs of the type removed; hybrid BGCs counted for both classes)

The 1,417 BGCs were then analysed with BiG-SLiCE’s query mode in order to calculate their distance (*d*) from a set of pre-computed gene cluster families (GCFs) comprised of 1.2 mio known BGCs. The analysis showed that 845 out of 1,417 BGCs (59.6%) had a *d* > 900, indicating that they were only distantly related to a GCF. Fifty-five outliers were found with *d* > 1800, indicating extremely divergent BGCs. A wide span of distances was present within each phylum which indicates that each phylum contained BGCs that are both closely and distantly related to known BGCs (Figure 2B). The median distances showed significant variation between phyla, with Bacteroidota containing the highest novelty (median *d* = 1227) and Planctomycetota the lowest (median *d* = 742). This overall score was, however, influenced by the fact that different classes of BGC scored differently. For example, NRPS/PKS BGCs scored higher than e.g. terpenes or bacteriocins. Rankings of single BGC classes showed that the high Bacteroidota score was partly driven by the large number of NRPS (Figure 2C) and the small number of terpenes and bacteriocins (Figures 2E and F) in the phylum. This is evidenced by the fact that other phyla scored the highest in individual BGC classes. For NRPS BGCs, Gemmatimonadota, Acidobacteriota and Verrucomicrobiota showed the highest values for *d* (Figure 2C). Gemmatimonadota furthermore showed the highest value for *d* when considering terpene BGCs (Figure 2E), while Acidobacteriota scored high for lassopeptides, arylpolyenes and PKS (Figure 2G,H,D). To check whether low coverage and the resulting insertion and deletion errors in the assembly led to overestimation of *d*, contig coverage as well as percentage of correctly-sized ORFs (as calculated by ideel) were plotted against *d*. There was no correlation between percentage of correctly sized ORFs and distance, indicating no effect of truncated ORFs on distance estimation. There was a slight positive correlation of *d* values with increased coverage, indicating a light, counterintuitive underestimation of novelty at low coverage. As expected, coverage showed a strong positive correlation with percentage of correctly-sized ORFs (see Supplementary Figures 1-3).

### Acidobacterial BGCs

Analysis of acidobacterial BGCs by order (Figure 3A) showed that terpenes were the most numerous, but with significant contributions from PKS, NRPS, lassopeptide and bacteriocin clusters. The orders of Pyrinomonadales and Vicinamibacterales constituted >60% of BGCs.

**Figure 3:**
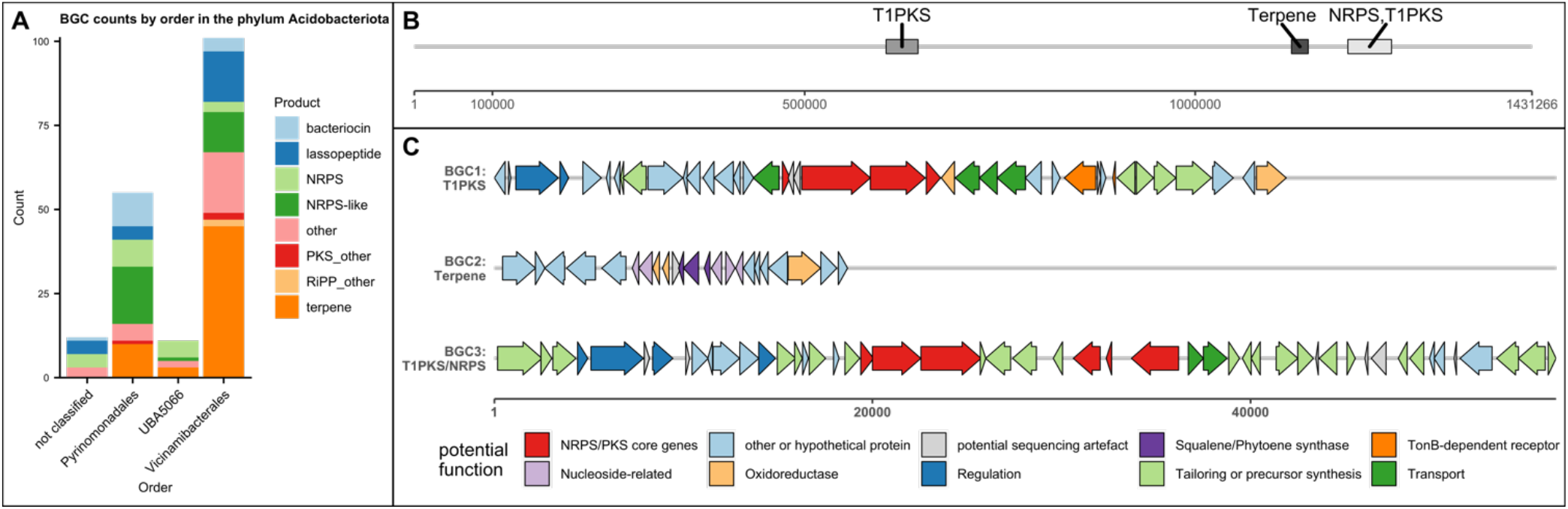
(A) BGC counts by BGC type and order in phylum Acidobacteriota; (B) Map of a large Acidobacteriota contig (order Vicinamibacterales) and the BGCs on it (C) Cluster map of proposed functions of genes in BGC1, BGC2 and BGC3. Functions were predicted from BLASTing against NCBI nr database as well as antiSMASH module predictions. A detailed table of homologous proteins can be found in the supplementary files

BiG-SCAPE analysis showed that BGCs mainly clustered together within orders (Supplementary Table 1). None of the families contained MiBiG clusters at the cut-offs used. Acidobacteriota showed a large number of lassopeptides, 16 of which grouped into two GCFs. NRPS-like BGCs also contributed a large number to the sample. In particular, one NRPS-like family from the order Vicinamibacterales showed homology to the VEPE BGC from *Myxococcus xanthus* in ClusterBlast. Furthermore, seven NRPS/PKS with a megasynthase gene length of over 20 Kb were found with the largest BGC measuring 89 Kb of NRPS and PKS megasynthase genes. The largest Acidobacteriota (order Vicinamibacterales) contig was 1.5 Mb in size and contained three BGCs: a PKS, a terpene and a NRPS/PKS hybrid cluster (Figure 3B,C). BGC1 (*d* = 1397) contained a partial one-module NRPS followed by a partial PKS module as well as transporter genes and a TonB-dependent receptor protein, suggesting a role as a siderophore. BGC2 (*d* = 1103) contained squalene/phytoene synthase genes and several potential tailoring enzymes. BGC3 (*d* = 1977) contained a complete NRPS and a partial NRPS module and an incomplete PKS domains. Several gaps visible in the BGC make a sequencing error seem possible, leading to truncated genes and therefore missing domains.

### Verrucomicrobial BGCs

The analysis of Verrucomicrobial BGCs by order (Figure 4A) showed that the vast majority of BGCs were terpenes, followed by arylpolyenes, PKS, NRPS, as well as ladderanes. The most prolific producer orders were Opitutales, Pedosphaerales and Chtoniobacterales.

**Figure 4:**
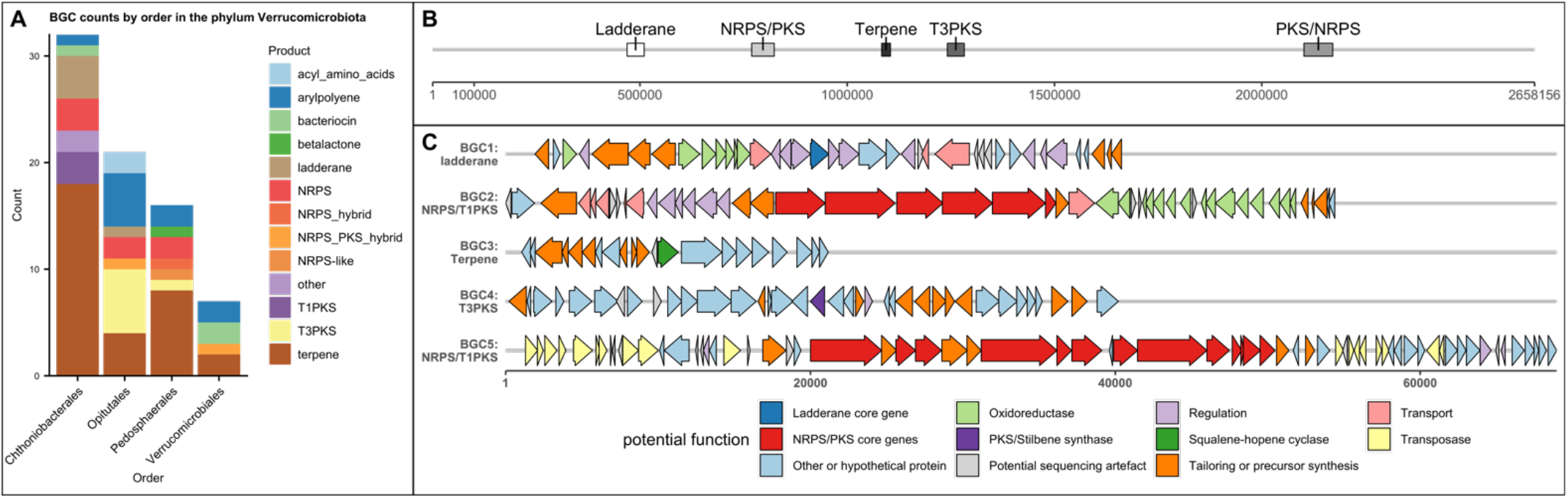
(a) BGC counts by BGC type and order in phylum Verrucomicrobiota, (b) map of a large Verrucomicrobiota contig (order Opitutales) and the BGCs on it; (b) Cluster map of proposed functions of genes in BGC1 – BGC5. Functions were predicted from BLASTing against NCBI nr database as well as antiSMASH module predictions. X axis represents basepairs. A detailed table of homologous proteins can be found in the supplementary files

Verrucomicrobial BGCs did not show strong clustering into conserved GCFs compared to Acidobacteriota (Supplementary Table 2). One NRPS and one PKS BGC were the only BGCs that clustered with MiBiG clusters.

The largest Verrucomicrobiota contig (order Opitutales) was 2.6 Mb in size and featured five BGCs, two of which were NRPS-PKS hybrids with megasynthase genes above 20 Kb (Figure 4B, C). BGC1 (*d* = 1479) contained a ladderane-type 3-oxoacyl-[acyl-carrier-protein] synthase. BGC2 (*d* = 1305) contained four NRPS modules interspersed by one PKS module. BGC3 (*d* = 673) contained a squalene-hopene cyclase, indicating a role in hopanoid biosynthesis. BGC4 (*d* = 1142) encoded a chalcone/stilbene synthase. BGC5 (*d* = 1340) contained a PKS module followed by five NRPS modules. The third module, however, showed a truncated A domain, with the antiSMASH HMM NRPS-A_a3 only matching around 50 bp at the end of ORF ctg423_1968. This could be explained by a sequencing error in which an indel lead to a frameshift, causing a premature stop codon. Indeed, nucleotide-level BLAST of the gap between ctg423_1968 and the PCP-domain containing ctg423_1970 showed a match to known A domains. It is, however, not possible to rule out potential pseudogenisation.

### Uncultivated and underexplored classes and orders from Actinobacteriota and Proteobacteria show a large biosynthetic potential

#### Actinobacteriota: Acidimicrobiia and Thermoleophilia

The phylum Actinobacteriota (335 BGCs) featured a large amount of BGCs unclassified at order level. Therefore, they were analysed by class (Figure 5A). The class Actinobacteria (114 BGCs) contained BGC-rich genera such as *Streptomyces* and *Pseudonocardia* and accordingly contributed a large amount of BGCs in the sample. The class Acidimicrobiia (90 BGCs) contained the genera *Illumatobacter* and *Microthrix* and several uncultured genera. The class Thermoleophilia (95 BGCs) contained genera such as *Solirubrobacter* and *Patulibacter*, besides uncultured genera, and contributed to a large amount of the bacteriocin and betalactone BGCs. The amount of BGCs in these classes that were not placed into lower taxonomic ranks indicated that there is a large unexplored diversity of uncultured Actinobacteriota containing a great diversity of BGCs.

**Figure 5:**
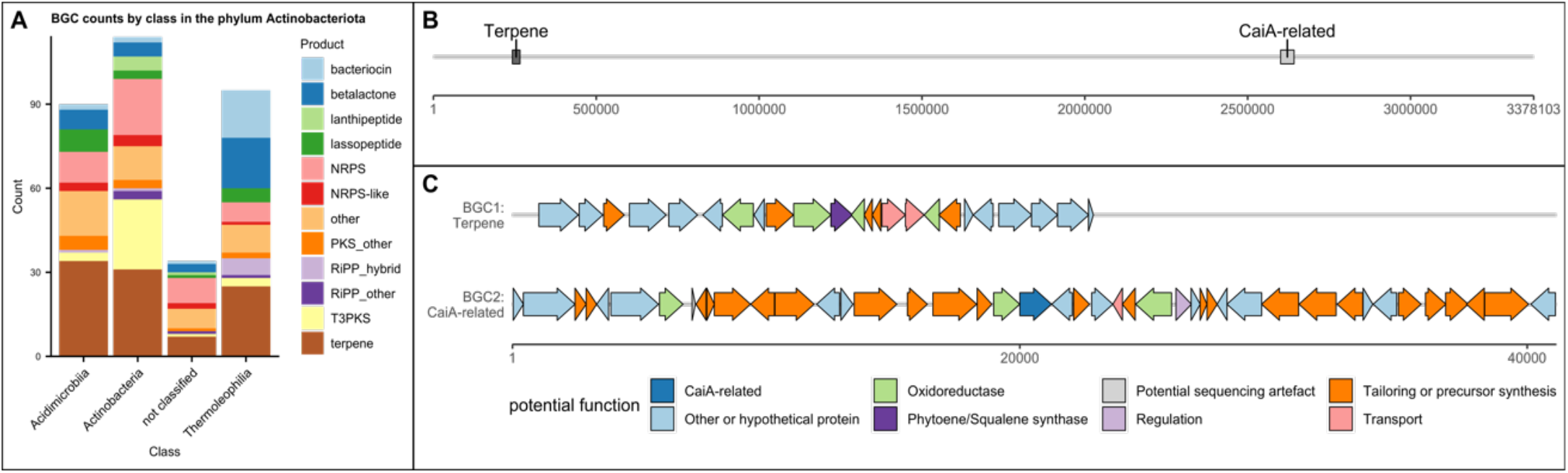
(a) BGC counts by BGC type and class in Actinobacteriota; (b) Map of a large Actinobacteriota contig (order IMCC26256) and number of basepairs; (c) Cluster map of proposed functions of genes in BGC1 and BGC2. Functions were predicted from BLASTing against NCBI nr database as well as antiSMASH module predictions. X axis represents basepairs. A detailed table of homologous proteins can be found in the supplementary files

Remarkably, one circular genome from the uncultured order IMCC26256 from the class Acidimicrobiia was recovered in a single contig, measuring 3.3 Mb in size and containing two BGCs (Figure 5B-C). The terpene BGC (*d* = 1398) contained a squalene synthase, a lycopene cyclase and polyprenyl synthetases, suggesting a role in pigment formation. The CaiA-related BGC (*d* = 1869) contained an acyl-CoA dehydrogenase related to CaiA (involved in saccharide antibiotic BGCs). BLAST hits indicated other genes related to small organic acids, sugars and nucleoside metabolism.

Two families of terpenes containing terpene cylases, methyltransferases and/or P450s showing similarity to the known geosmin and 2-methylisoborneol BGCs were found, with members belonging to both Acidimicrobiia, Thermoleophilia and unclassified Actinobacteriota. One BGC from a *Streptomyces* spp. was detected, containing a LmbU-like gene on the very edge of the contig. BiG-SCAPE analysis showed that Actinobacteriota BGCs mostly grouped within the classes, and one lanthipeptide BGC grouped with MiBiG BGCs at the cut-off used (Supplementary Table 3).

### Proteobacteria: the uncultured methanotrophic order UBA7966 as a specialised metabolite producer

Analysis at the order level of the proteobacterial BGCs showed that the biggest contributor was the Burkholderiales order with 116 BGCs (Figure 6A) followed by order UBA7966 with 96 BGCs. UBA7966 BGCs included a variety BGCs, including terpenes, bacteriocins, phosphonates, NRPS & NRPS hybrids, NRPS-like, and arylpolyenes. In particular, the high abundance of NRPS-like and phosphonate BGCs in UBA7966 contrasted with the lower counts in other proteobacterial orders in the dataset. By order, UBA7966 contigs also showed a high average coverage 26x, compared to the total average of 10.2x, indicating a high abundance. The total length of UBA7966 contigs was 53 Mb, indicating the presence of several genomes.

**Figure 6:**
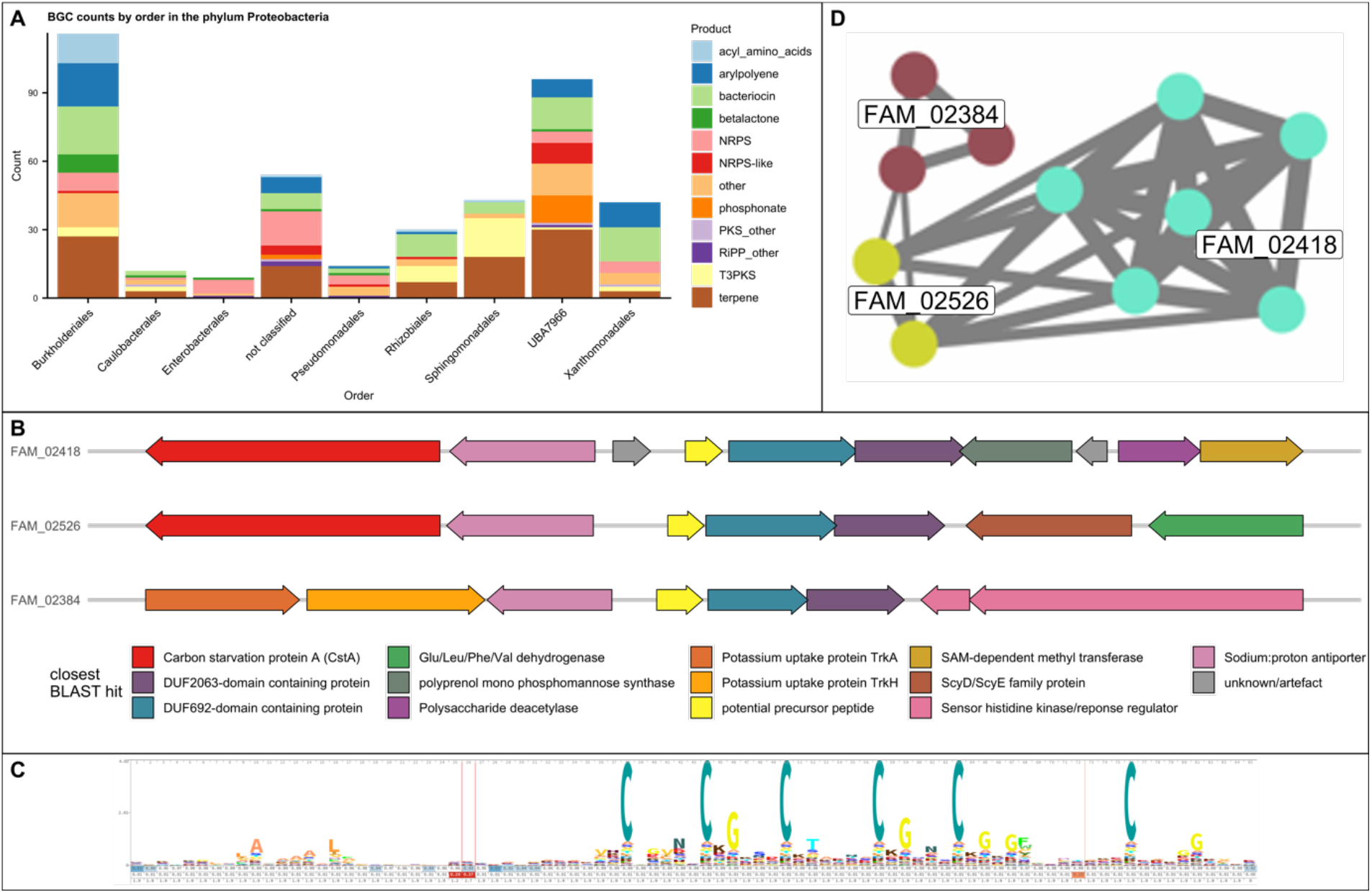
(a) BGC counts by BGC type and order in the phylum Proteobacteria; (b) Cluster layout of three gammaproteobacterial DUF692-containing BGCs representatives: contig_12391 for FAM_02418, contig_14956 for FAM_02526, and scaffold_15362 for FAM_02384; (c) Sequence logo generated from an HMM of 301 potential precursor peptides; (d) Similarity network generated from BiG-SCAPE with brown: FAM_02384, turquoise: FAM_02418, green: FAM_02526.

The order UBA7966 is an uncultured order consisting of one family, UBA7966, which contains two genera, *UBA7966* and *USCγ-Taylor*. UBA7966-family bin bin.3 was assigned no genus, while all CAT-assigned contigs were assigned species *USCγ-Taylor sp002007425*, the only species in the *USCγ-Taylor* genus. The *USCγ-Taylor* genus is based on a putatively methanotrophic MAG extracted from a methane-oxidising soil metagenome from Taylor Valley in Antarctica (Genbank accession GCA_002007425.1)^26^. The low number of UBA7966 reference genomes in the GTDB database means, however, that these classifications remain an approximation. The two closest orders to UBA7966 that contain cultured representatives, Beggiatoales and Nitrosococcales, both have members implicated in methanotrophy, sulphur cycling and ammonia oxidation as well as varying degrees of chemolithotrophy and chemoautotrophy^46–49^. On all the contigs assigned to order UBA7966 by CAT, four *pmoCAB* operons were found, with *pmoA* showing 92.9% to 96.8% identity with *pmoA* from *USCγ-Taylor*. This indicates that, in addition to the methanotrophy of *USCγ-Taylor*, other members of the order UBA7966 could be involved in similar lifestyles.

When analysed with BiG-SCAPE at cut-off 0.7 (Supplementary Table 4), phosphonates (median *d* = 1421), NRPS/NRPS-like (median *d* = 1262) and bacteriocins seemed to form especially conserved GCFs. Other GCFs were shared with other proteobacterial orders. With 96 BGCs, UBA7966 contributed a similar number of BGCs as the established specialised metabolite producing order Burkholderiales (116 BGCs). However, the BiG-SLiCE distances of UBA7966 were higher than Burkholderiales for all but one BGC class, indicating more novel BGCs (Supplementary Figure 4).

The potential methanotrophy of UBA7966 suggested the potential presence of methanobactins, but no BGCs corresponding to known methanobactins were found in the dataset. On the other hand, an abundance of DUF692-containing BGCs were observed, grouping into three GCFs. DUF692 proteins are a diverse family of proteins with largely unknown functions, although some are known to be involved in methanobactin biosynthesis^50^. The analysis of three related GCFs containing DUF692 domains (including BGCs from UBA7966 and unclassified gammaproteobacterial contigs) showed that FAM_02526 (two BGCs), FAM_02384 (three BGCs) and FAM_02418 (six BGCs) (Figures 6B and D) all contained a short (circa 240 bp) ORF followed by first a DUF692-domain containing protein, then a DUF2063-domain containing protein. Furthermore, a putative cation antiporter was found upstream of the precursor peptide. The three families differed by the genes surrounding this core cluster (Figure 6B). The 11 small translated 240bp ORFs were aligned using Clustal Omega and a HMM search was made in ebi reference proteome database with a cut-off E-value of 10E-10. The resulting 290 protein sequences (almost exclusively from Proteobacteria) plus 11 original sequences were aligned using Clustal Omega and a HMM was generated and visualised using skylign.org. The resulting logo showed a low degree of sequence conservation except for a pattern of six conserved cysteines – some followed by glycines – within forty amino acids towards the N-terminus, and a slightly conserved hydrophobic patch towards the C-terminus (Figure 6C). This might represent a potential precursor peptide, with the six cysteines marking the potential core peptide.

The UBA7966 order also contained larger BGCs such as four NRPS/ NRPS-PKS BGCs with megasynthase genes with a length of more than 20 Kb, the largest cluster possessing 56 Kb of PKS (seven modules) along with NRPS (three modules) genes. This latter BGC also formed a BiG-SCAPE GCF with several MiBiG BGCs which shared the presence of a small peptide moiety followed by several malonyl units.

### Low numbers of BGC found in other underexplored phyla

Lower numbers of biosynthetic gene clusters were detected in the phyla Gemmatimonadota (31 BGCs), Planctomycetota (29), Myxococcota (22), Patescibacteria (9), Methylomirabilota (5), Bdellovibrionota_B (8), Elusimicrobiota (4), Armatimonadota (4) and Binatota (3) (Figure 7A).

**Figure 3:**
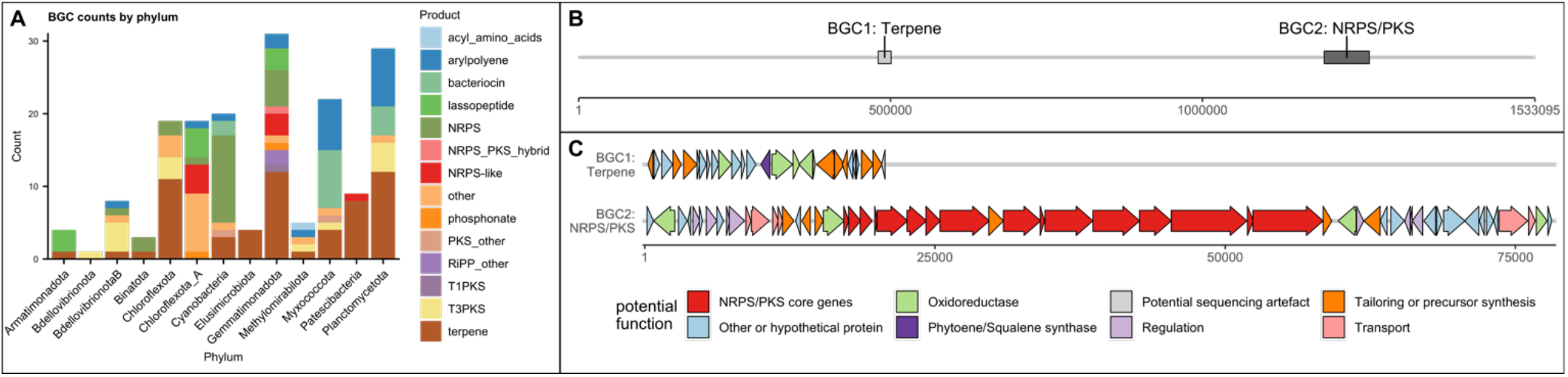
(A) Distribution of BGCs among phyla with 31 or fewer BGCs in the dataset; (B) Map of a large Gemmatimonadota contig (order Gemmatimonadales) and BGCs detected on it; (C) Cluster map of proposed functions of genes in BGC1 and BGC2. Functions were predicted from BLASTing against NCBI nr database as well as antiSMASH module predictions. X axis represents basepairs. A detailed table of homologous proteins can be found in the supplementary files

One remarkably long (1.5 Mb, Figure 7B,C) Gemmatimonadota contig from the order Gemmatimonadales was found to contain two BGCs: one terpene (*d* = 998) and one NRPS/PKS BGC (*d* = 1423). BGC1 contained a phytoene synthase and several related oxidases. BGC2 contained six PKS modules and two NRPS modules as well as modifying enzymes presence of a TonB receptor indicated that the product could serve as a siderophore.

## Discussion

### Metagenomics reveal biosynthetic potential of underexplored bacterial lineages

In our dataset, we found a large number of BGCs in underexplored phyla not usually associated with specialised metabolites. Two previous studies noted NRPS and PKS novelty and diversity in Acidobacteria and Verrucomicrobia^7,8^. The present study indicates that these underexplored phyla harbour not only novel NRPS/PKS, but new BGCs from many different classes, such as lassopeptides and bacteriocins. While Crits-Cristof et al. ^7^ highlighted two promising acidobacterial MAGs from the classes Blastocatellia and the Acidobacteriales, in the present sample the classes Blastocatellia and Vicinamibacteria were the main contributors of acidobacterial BGCs. Furthermore, many BGCs were found in other ubiquitous phyla such as Patescibacteria, Gemmatimonadota and Armatimonadota. Three BGCs (two NRPS and one terpene) were placed in the phylum Binatota. The phylum Binatota was proposed by Chuvochina et al. based on a handful of soil MAGs with no cultured representatives^37^. To our knowledge, this is the first description of BGCs belonging to the phylum Binatota. We also discovered highly divergent BGCs from the underexplored Actinobacteriota classes Acidimicrobiia and Thermoleophilia. This indicates that Actinobacteriota, which contain the heavily exploited genus *Streptomyces*, contain unknown lineages harbouring interesting BGC diversity.

In the present dataset, 845 out of 1,417 BGCs (59.6%) had a *d* > 900 and 55 (3.9%) had a *d* > 1800 to the closest GCF. These numbers contrast starkly with the 1.2 million original BGCs in the BiG-SLiCE dataset, of which only 13.9% and 0.2% showed *d* > 900 and *d* > 1800 respectively. While it is necessary to note that sequence diversity does not demonstrate chemical diversity, the striking amount of sequence divergence encountered in just one soil sample adds to the mounting evidence that uncultured and underexplored phyla – especially Acidobacteriota – are promising candidates for the discovery of novel specialised metabolites. It is furthermore worth noting that the great biosynthetic diversity found at Mars Oasis is under threat from climate change, with the maritime Antarctic warming by 1– 3 °C between the 1950s and the turn of the millenium^51^, and, despite a recent pause in this warming trend^52^, similar increases in temperature being predicted for later this century as greenhouse gases continue to accumulate in the atmosphere^52,53^.

The large number of terpene BGCs, most of them putatively C30/C40 carotenoids or hopanoids, could be interpreted with respect to the roles of these compounds in membrane function at extreme temperatures^22,54,55^, as well as UV protection^22,56^. A previous study similarly noted a high number of pigmented bacteria among isolates from Antarctic samples^22^. Kautsar et al.^42^ recorded only 7.8% terpene BGCs in their large-scale survey of publicly available bacterial genomes, as opposed to the ca. 25% in this survey. Previous short-read metagenomic studies of aquatic and soil environments also reported high numbers of terpene BGCs, with terpenes representing between 15% and 50% of the reported BGCs, respectively^57–59^. However, the representativeness of BGC counts obtained through metagenomic studies remains questionable. Small terpene BGCs are easier to assemble than long and repetitive NRPS/PKS BGCs, therefore leading to bias.

In this study, a large number of BGCs were observed in potentially methanotrophic members of the uncultured order UBA7966. Methanotrophic organisms have not usually been linked to specialised metabolite production, except for siderophore-like RiPPs called methanobactins able to scavenge the copper needed for methane and/or ammonia oxygenase enzymes^50^. We reason that the lack of known natural products might be related to difficulties associated with cultivation such as specific nutrient requirements and often slow growth, as well as to the amount of energy, carbon and nitrogen available for specialised metabolite production. While no methanobactin BGCs were seen in UBA7966-classified contigs, examining three gammaproteobacterial DUF692-domain containing GCFs revealed the presence of a potential conserved six cysteine precursor peptide. The conserved cysteines in the potential precursor peptides are resemblant of ranthipeptides (formerly known as SCIFFs), which contain six cysteines in forty-five amino acids. Ranthipeptides, however, contain thioethers formed by radical SAM enzymes^60^. DUF692 domain proteins are furthermore known to be involved in methanobactin and TglA-thiaGlu biosynthesis^50,61^, and at least one member has been shown to contain two iron atoms potentially acting as cofactors^61^. All DUF692 protein containing GCFs in the order UBA7966 observed in the present study also contained DUF2063 proteins. DUF2063 family proteins are mostly uncharacterised, though the crystal structure of a member from *Neisseria gonorrhoeae* indicates that DUF2063 might be a DNA-binding domain involved in virulence, and there has been one report of co-occurrence of DUF2063 and DUF692 proteins^62^. Other studies discovered the two neighbouring proteins in operons related to stress response at high calcium concentration^63^ in *Pseudomonas* as well as responding to gold and copper ions^64^ in *Legionella*. The two genes were also found in the atmospheric methane oxidiser *Methylocapsa gorgona*^65^. We therefore hypothesize that these BGCs could be another form of RiPP involved in chelating metals. While the six cysteines could be involved in forming thioether bonds, disulfide bonds or lanthithionine groups like in many other RiPPs, they could potentially also be directly involved in metal coordination as is the case in the group of small metal-binding proteins called metallothioneins^66^.

### Long reads make mining and phylogenetic classification of metagenomic BGCs feasible

The advantage of long reads could be observed from comparing the results achieved from long reads vs. short reads, with the short reads providing a lower number of BGCs and a significantly lower taxonomic classification success compared to the BGCs assembled and annotated using long reads. While the number of bases used in the assembly was about a third lower for short reads (28 Gb vs 44 Gb), the number of recovered BGCs was more than two thirds lower (430 BGCs vs 1,417 BGCs) and the BGCs assembled from short reads were mostly incomplete. Moreover, this study showed that long-read metagenomes constitute a valuable tool to achieve similar or even improved results to previously very expensive deep short-reads metagenomes ^7,57,58^. For example, Cuadrat et al. used 500 million reads (*c*. 50 Gb if read length was 100 bp) for BGC genome mining of a lake community recovering 243 BGCs with a total of 2,200 ORFs, which averages to nine ORFs per BGC indicating small and/or incomplete BGCs^58^. A larger short-read study of microbial mats recovered 1,477 BGCs^57^. While this study did not report the number of sequenced bases or BGC completeness, the median BGC length of 103 BGCs from 15 representative and highly complete MAGs was 11.9 Kb, also indicating mostly small and/or incomplete BGCs. Another study by Crits-Cristof et al. ^7^ used 1.3 Tb of short-read sequence data of grassland soil to mine selected bins of four phyla, recovering a total of 1,599 BGCs, 240 of which were NRPS/PKS BGCs, including several large and complete ones^7^. The present study indicates that the long-read approach requires a relatively low sequencing input similar to the two smaller studies to provide a result similar to the larger study. While the contigs, MAGs and BGCs produced using shallow ONT sequencing are not as accurate as the ones produced using deep short read sequencing, our results show that they are sufficiently accurate to profile the biosynthetic potential of complex environmental samples, estimate their diversity and could be used to guide isolation and heterologous expression strategies. Lower error rates could be achieved through higher coverage in long and short reads as well as advances in long-read basecalling. We furthermore conclude that contig-level classification using CAT shows advantages compared to genome-resolved metagenomics in single-sample data, where binning is inefficient. Cuadrat et al, Crits-Cristof et al. and Chen et al. used genome-resolved metagenomics^7,57,58^, in which contigs are binned and bins are mined for BGCs. While it is favourable to attribute BGCs to distinct MAGs, it is viable only when a large number of samples are used, making binning efficient through differential abundance^67^. When using only one sample, binning becomes inefficient and, in our case, missed the vast number of BGCs, with 1,139 of 1,417 BGCs not being binned. Contig-based classification approaches offer an alternative, but their accuracy is limited by contig length^35^ and the classification dependent on the database used. In our data, a contig N50 of >80 Kb provided ample sequence data for accurate classification, leading to >90% classification at phylum level. Usage of GTDB-derived databases ensured improved classification of uncultured taxa, and few conflicts with single-copy core gene-based bin-level classification were detected.

## Conclusions and Perspectives

The use of nanopore metagenomic sequencing, binning and contig-based classification approaches using GTDB combined with BGC genome mining allowed us to identify 1,417 BGCs, 75% of which were complete, from a wide range of soil bacteria. This confirms and further expands our knowledge of the biosynthetic potential of difficult-to-culture phyla such as Verrucomicrobiota, Acidobacteriota and Gemmatimonadota. In addition, we show that uncultured and underexplored lineages of the well-known producer phyla Actinobacteriota (classes Thermoleophilia and Acidimicrobiia) and Proteobacteria (order UBA7966) show a large biosynthetic potential.

We furthermore demonstrate that ONT long-read sequencing enables the assembly, detection and taxonomic classification of full-length BGCs on large contigs from a highly complex environment using only one sample and <100 Gb sequencing data, which presents a >10-fold reduction compared to studies using short reads to recover large and complete BGCs. While more samples would be needed for improved binning and genome-resolved metagenomics, our approach proved successful in classifying 70% of BGCs at order level.

Even with limited sequencing, we were able to retrieve megabase-sized contigs and one circular genome containing multiple BGCs. With nanopore sequencing becoming more widespread, it will soon be commonplace to profile the biosynthetic potential of uncultured microbes from diverse environments without enormous sequencing efforts. In combination with heterologous expression techniques such as DiPaC^68^, accessing natural products from metagenomes could be revolutionised, overcoming the need for constructing, maintaining and screening large metagenomic libraries or large sequencing budgets. For remote and endangered environments such as the Antarctic Peninsula, which is warming rapidly due to climate change, these metagenomic strategies will prove especially valuable.

## Data availability statement

The nanopore and Illumina reads generated in this study have been deposited in the Sequence Read Archive with the accession code PRJNA681475 (https://www.ncbi.nlm.nih.gov/sra/PRJNA681475).

## Supporting information

BGC data

