## Supplemental Tables and Figures for "Metabolic potential of uncultured Antarctic soil bacteria revealed through long-read metagenomic sequencing"

### Supplementary Tables

Supplementary Table 1: GCFs with >2 members, at least one of which is a member of the Acidobacteriota phylum

| Family | No. BGCs | Order/s | BGC type | Notes |
| --- | --- | --- | --- | --- |
| 3163 | 6 | Not classified (5), Pyrinomonadales (1) | NRPS | A group of similar NRPS in various states of fragmentation featuring polysaccharide biosynthesis enzymes. |
| 1862 | 4 | Pyrinomonadales | NRPS-like | Similar to VEPE |
| 2375 | 4 | Vicinamibacterales | NRPS-like | All contain short chain dehydrogenases. NRPS-like domains: Adenylation + PP-attachment |
| 2575 | 3 | Pyrinomonadales | NRPS-like | All contain short chain dehydrogenases. NRPS-like domains: Adenylation + PP-attachment |
| 2590 | 7 | Pyrinomonadales | Terpene | Contain squalene/phytoene synthase |
| 2666 | 13 | Vicinamibacterales | Terpene/NRPS-like | Contain squalene/phytoene synthase |
| 3035 | 6 | Vicinamibacterales | Terpene | Contain squalene/phytoene synthase |
| 3174 | 3 | Vicinamibacterales | Terpene | Contain squalene/hopene cyclase |
| 2454 | 3 | Pyrinomonadales | T3PKS | Contain chalcone/Stilbene synthases |
| 1915 | 10 | Vicinamibacterales,  [Proteobacteria:] Rhizobiales, Burkholderiales, Caulobacterales, Sphingomonadales | Bacteriocin | DUF692-containing. Contain Sigma 70 2& 4 domains as well as different transmembrane proteins. Potentially related to methanobactins. |
| 1917 | 9 | Vicinamibacterales/ not classified | lassopeptide | All contain ABC transporters that have similarity to paeninodin lassopeptide BGC, some contain SAM-dependent methyltransferases that show similarity to DivMT from Divamide Lassopeptide. No precursor peptide predicted |
| 2119 | 3 | Pyrinomonadales | Bacteriocin | Radical SAM pf04055, containing a serine protease. Very small BGCs as annotated by antiSMASH. |
| 2238 | 7 | Pyrinomonadales | lassopeptide | All contain a conserved pattern of methyltransferases and glycosyltransferases. Looks like potential peptidoglycan biosynthesis enzymes, since phospho-N-acetylmuramoyl-pentapeptide-transferase is annotated. Precursor peptides annotated. |

Supplementary Table 2: GCFs with >2 members, at least one of which is a member of the Verrucomicrobiota phylum

| Family | No. BGCs | Order/s | BGC type | Notes |
| --- | --- | --- | --- | --- |
| 1967 | 9 | Pedosphaerales, [Proteobacteria:] Burkholderiales, [Gemmatimondota:] Gemmatimonadales, unclassified, | NRPS | This GCF family contains lots of NRPS that do not share any obviously discernable features. Contains BGC0001813.1 Tyrobetaine. |
| 2031 | 4 | Chthoniobacterales | Terpene | Contain squalene/phytoene synthases |
| 1255 | 9 | 2x Chthoniobacterales, rest mibig | T1PKS | Contains fungal MiBiG BGCs such as Pyranonigrin E, Terreic acid, Ochratoxin A, but does not show obvious similarities. |
| 3150 | 3 | Chthoniobacterales | ladderane |  |
| 3167 | 5 | Opitutales | Arylpolyene |  |

Supplementary Table 3: GCFs with >2 members, at least one of which is a member of classes Acidimicrobiia or Thermoleophilia

| Family | No. BGCs | Class/es | BGC type | Notes |
| --- | --- | --- | --- | --- |
| 1927 | 4 | Acidimicrobiia / not classified | NRPS-like, betalactone | NRPS-like and betalactone BGCs, with transporters |
| 1870 | 3 | Acidimicrobiia | Lassopeptide | Contain nucleodityltransferases, glycosyltransferases and ABC transporters |
| 2650 | 9 | Thermoleophilia | betalactone | Contain HMGL-like and AMP-binding domains |
| 2170 | 5 | Thermoleophilia | other | CaiA flavin-dependent oxidoreductase BGC. |
| 1893 | 3 | Thermoleophilia | betalactone | Contain HMGL-like and AMP-binding domains |
| 507 | 6 | 1x Thermoleophilia, rest mibig | Lanthipeptide/bacteriocin (DUF692) | Lanthipeptide part of BGC groups with mibig lanthipeptides. |
| 2409 | 4 | Thermoleophilia | bacteriocin | DUF692, all containing carbohydrate kinases, 3 contain transmembrane transporter |
| 2709 | 4 | 2x Thermoleophilia, 1x UBA7966 [Proteobacteria], 1x unclassified Actinobacteriota | bacteriocin | PF04454 |
| 2386 | 4 | Thermoleophilia Acidimicrobiia, C 2x unclassified ACtinobacteriota | Terpene | All show similarity to geosmin/2-methylisoborneol |

Supplementary Table 4: GCFs with >2 members, at least one of which is a member of the UBA7966 order

| Family | No. BGCs | Class/es | BGC type | BGC family features |
| --- | --- | --- | --- | --- |
| 1002 | 10 | 2x UBA7966, 2x unclassified, 6x MiBiG | NRPS, NRPS/T1PKS | Cluster with 6 Cyanobacteria-related BGCs such as Cyanopeptolin, Anabaenopeptin. |
| 1703 | 6 | 1x UBA7966, 5x mibig | NRPS, T1PKS | 5 Myxococcus-related BGCs like Myxalamid, Phenalamid, and Sorangium-related Pellasoren |
| 2200 | 5 | UBA7966 | NRPS-like | Feature Penicillin binding protein transpeptidase and transglycolase domains |
| 2458 | 4 | UBA7966 | NRPS-like | Similar to FAM_2020. Feature Penicillin binding protein transpeptidase and transglycolase domains (peptidoglycan biosynthesis) |
| 926 | 7 | 1x UBA7966, 1x Burkholderiales, 5x mibig | siderophore | Clusters with desferrioxamins |
| 1901 | 5 | UBA7966, 1x unclassified | Phosphonate | All similar to Metcalf et al Group 1: Burkholderia-like phosphonolipids |
| 2294 | 4 | UBA7966 | Phosphonate | All featuring an aminotransferase class V andor a lysylphosphatidylglycerol synthase TM region, similar to Group 2 from Metcalf et al. |
| 2367 | 5 | UBA7966, 1x Burkhoderiales | Arylpolyene, Arylpolyene/terpene |  |
| 2525 | 5 | UBA7966, 1x unclassified | Phosphonate | All featuring an aminotransferase class V and/or a lysylphosphatidylglycerol synthase TM region, similar to Group 2 from Metcalf et al. |
| 2743 | 4 | UBA7966, 1x Xanthomonadales | Arylpolyene, Arylpolyene/terpene |  |
| 2807 | 5 | 1x UBA7966, 1x Xanthomonadales, 3x unclassified | Arylpolyene |  |
| 2941 | 7 | UBA7966, 1x unclassified | Arylpolyene, Arylpolyene/terpene |  |
| 2384 | 3 | 1x UBA7966, 2x not classified | Bacteriocin | DUF692 with cation transporter |
| 2418 | 6 | UBA7966 | Bacteriocin | DUF692 with CstA transporter and sodium/calcium exchanger, very conserved synteny |
| 2709 | 4 | 1x, UBA7966, 2x Solirubrobacterales, 1x unclassified Actinobacteriota | Bacteriocin | PF04454 |
| 2047 | 11 | UBA7966 | Terpene | Squalene-hopene cyclase and phytoene synthase |
| 2518 | 13 | UBA7966 | Terpene | Phytoene synthase |
| 2574 | 3 | UBA7966 | Terpene | Phytoene synthase |

Supplementary Table 5: GCFs with >2 members, at least one of which belonging to the phyla Gemmatimonadota, Planctomycetota, Myxococcota, Patescibacteria, Methylomirabilota, Bdellovibrionota_B, Elusimicrobiota, Armatimonadota or Binatota

| Family | No. BGCs | Phylum | BGC type | BGC family features |
| --- | --- | --- | --- | --- |
| 1848 | 3 | Armatimonadota | Lassopeptide | Contain citydylyltransferase and DNA topoisomerase domains |
| 2244 | 4 | Patescibacteria | Terpene | Contain squalene/phytoene synthase |
| 2937 | 3 | Gemmatimondatoa | Lassopeptide | Contain ABC transporter and LuxR regulator |
| 1967 | 9 | Gemmatimonadota, Verrucomicrobia, Proteobacteria, unclassified | NRPS | This GCF family contains several NRPS/PKS from different lineages, Contains BGC0001813.1 Tyrobetaine. |
| 2373 | 4 | 3x Gemmatimonadota, 1x unclassified | Terpene | squalene/phytoene and carotenoid synthesis protein |

### Supplementary Figures

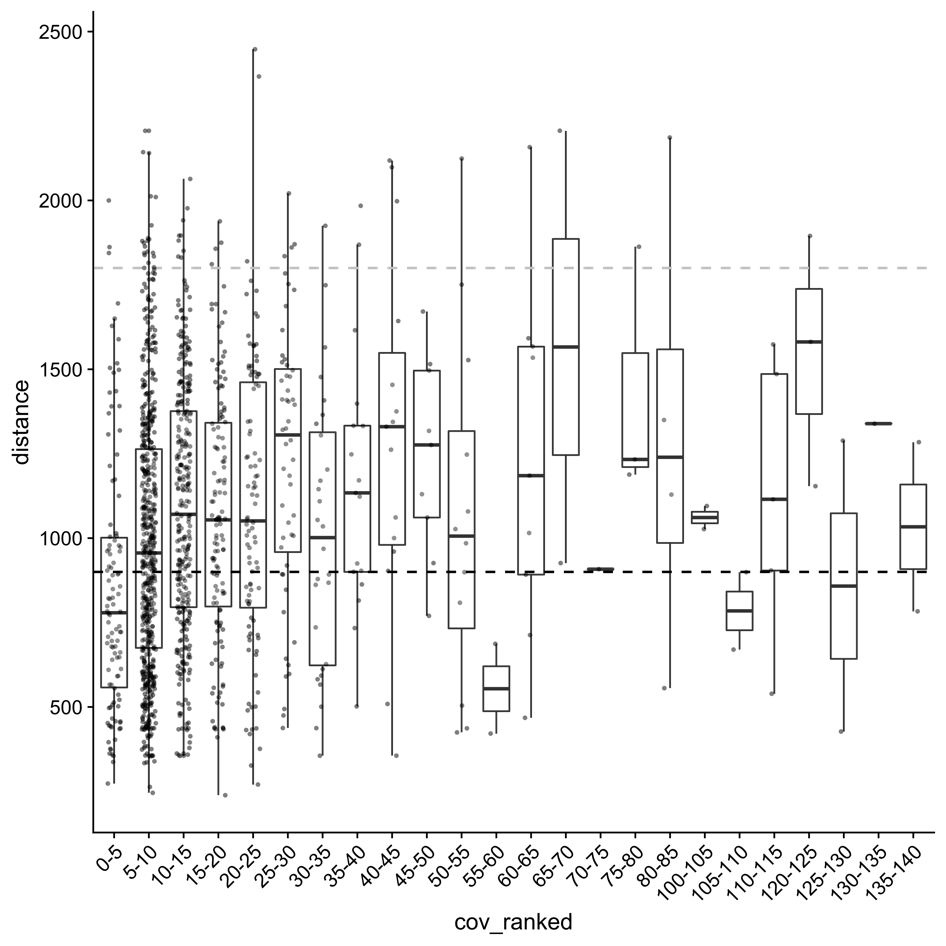

Supplementary Figure 1: Ranked coverage compared to BiG-SLiCE distance. Each point indicates a data point. A slight trend between coverage and distance is visible, especially between Coverage 0-5 to 10-15.

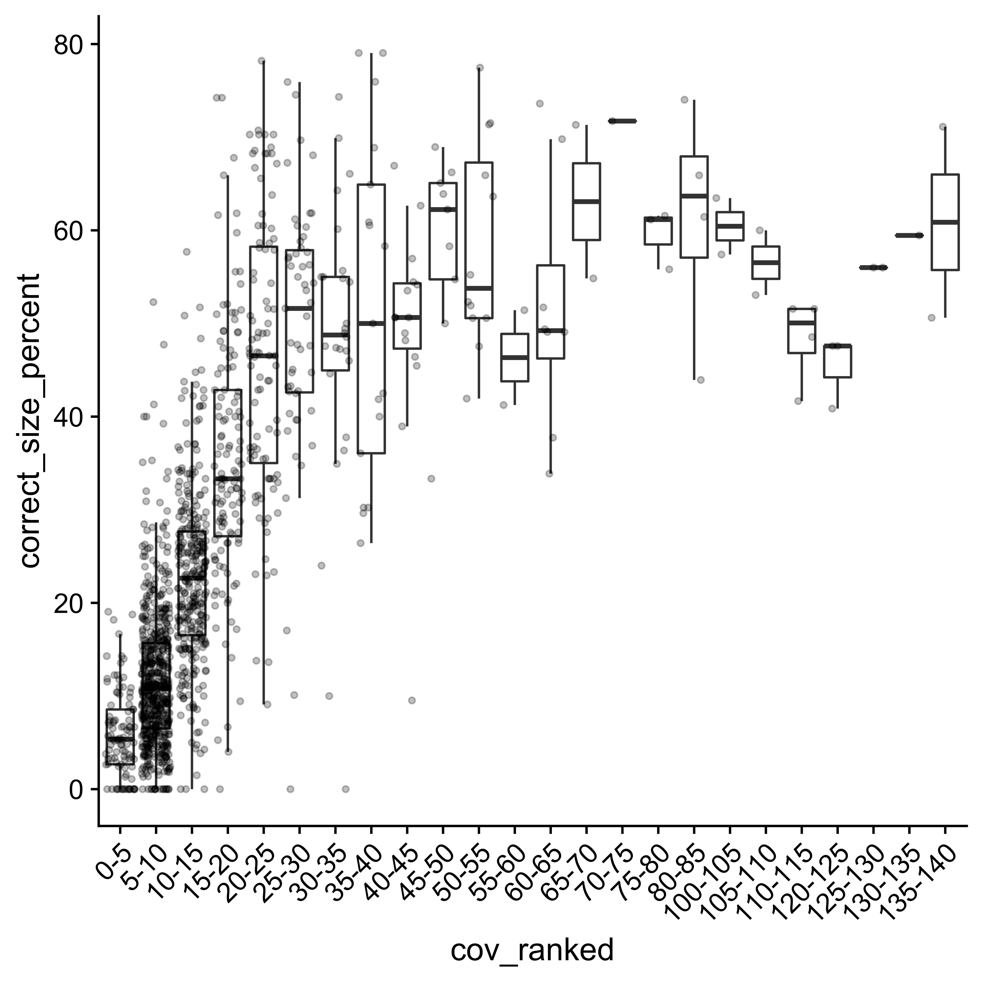

Supplementary Figure 2: Ranked coverage of BGC-containing contigs compared to percentage of “correct size” ORFs on same contig. “correct size” is defined by being between 0.9 to 1.1-times the length of a reference protein as calculated by ideel. Each point indicates a data point. A strong trend between coverage and number of “correct size” ORFs is visible up until coverage 25-30.

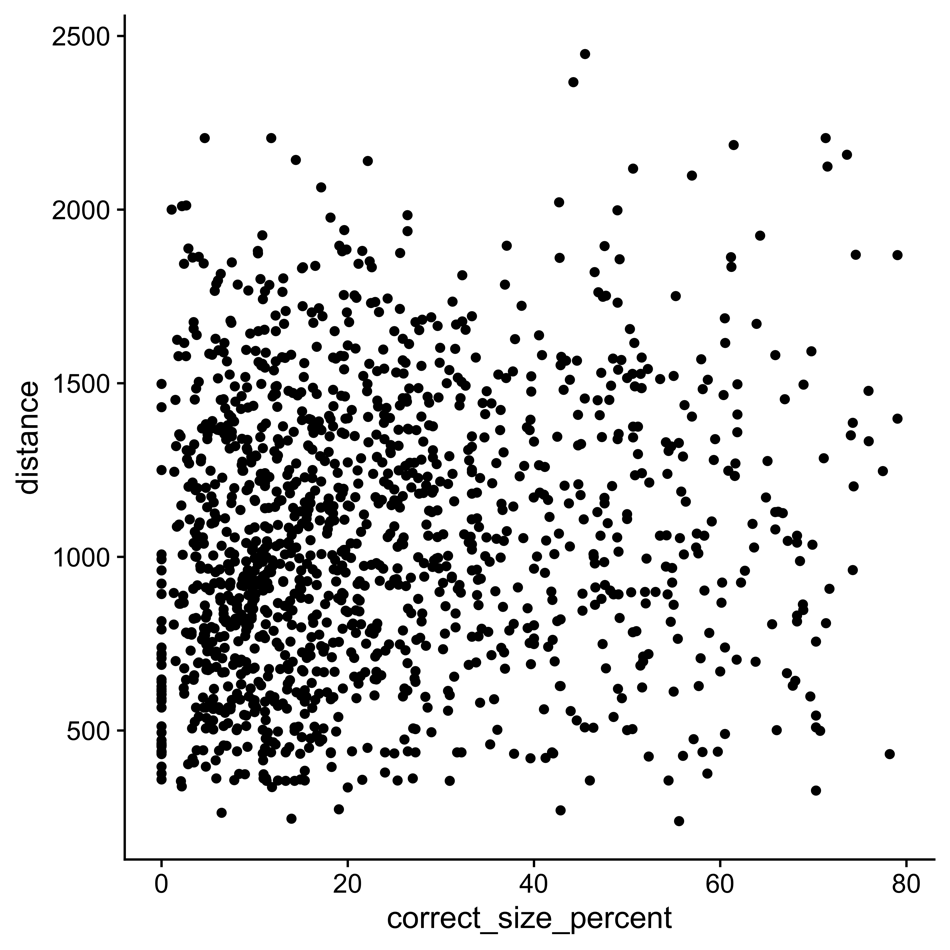

Supplementary Figure 3: Percentage of “correct size” ORFs on a BGC-containing contig compared to bigslice distance of the BGC. “correct size” is defined by being between 0.9 to 1.1-times the length of a reference protein as calculated by ideel. Each point indicates a data point. No correlation can be observed..

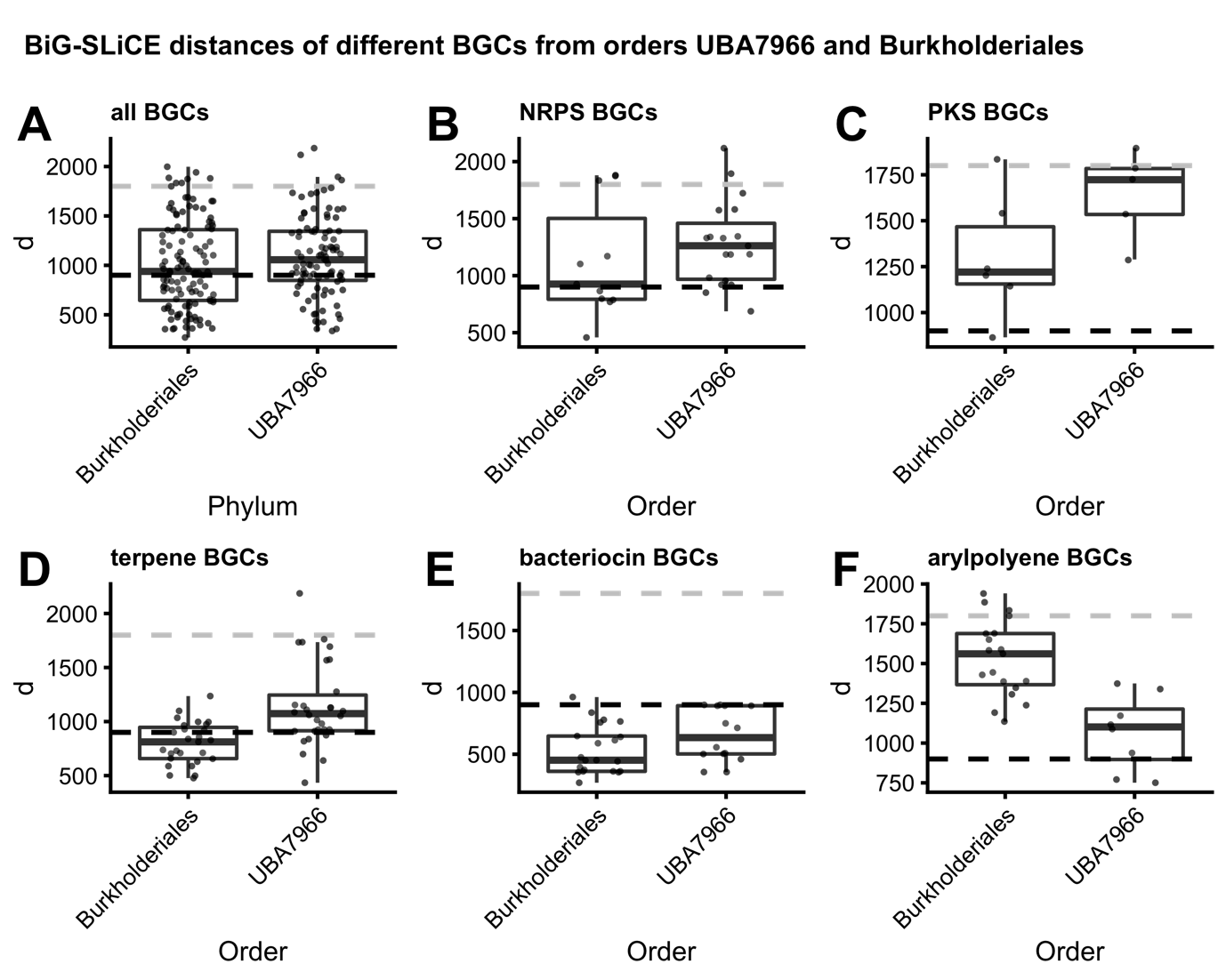

Supplementary Figure 4: Comparison of distances of BGCs of different classes between UBA7966 and Burkholderiales orders. (A) all BGCs; (B-F) specific BGC classes.
